# Divergent use of metabolic fluxes in breast cancer metastasis

**DOI:** 10.1101/2020.08.03.234468

**Authors:** Deepti Mathur, Chen Liao, Alessandro La Ferlita, Salvatore Alaimo, Alfredo Ferro, Joao B. Xavier

## Abstract

Breast cancers can metastasize to many organs. But how do disseminated cells from a primary tumor adapt to distal tissues? Here we combined metabolomics, flux measurements, and mathematical modeling to study metabolic fluxes in breast cancer cells adapted to home to different organs. We found that lung-homing cells maintain high glycolytic flux despite low levels of glycolytic intermediates, by constitutively activating a pathway sink into lactate. Their distinct behavior—a strong Warburg effect—has a gene expression signature: a high ratio of lactate dehydrogenase to pyruvate dehydrogenase gene expression, which also correlates with lung metastases in patients with breast cancer. Surprisingly, this strong Warburg effect does not necessarily increase cellular growth rate, suggesting that lactate secretion may be a trait under selection in lung metastasis. Our results stress that metabolic fluxes may not correlate with metabolic intermediates, a finding relevant for metastatic tropism.

## Introduction

Breast cancer is a disease marked by cellular diversity. Cancer cells differ at their phenotypic, genetic and proteomic levels both between and within patients (Turashvili and Brogi 2017). Cells in metastases can differ from cells in the primary tumor and even from those in metastases found in different organs (Turashvili and Brogi 2017; Minn, Kang, et al. 2005). Understanding the molecular mechanisms underlying such cellular diversity is vital for the future of targeted therapy.

Metastasis is a rare and stochastic process. For a breast cancer cell to form a metastasis it must invade the tissue that surrounds the primary tumor, intravasate into blood vessels, survive bloodstream circulation, extravasate from blood vessels at a distant site, and finally transition from a micro-to a macro-metastasis at the distant site, often after a period of dormancy that can last years. Each of these steps is inefficient and stochastic, and because of that it becomes very difficult to predict when, where and whether a patient diagnosed with breast cancer will develop metastases and which organs will be affected (Quail and Joyce 2013).

When a cancer cell disseminates from a breast tumor it may already have a propensity to metastasize to a specific organ (Minn, Gupta, et al. 2005; Nguyen et al. 2009; Minn, Kang, et al. 2005). The possibility that the distal tissue selects for specific features of cancer cells was first raised by Paget and is called the “seed-and-soil hypothesis” (Paget 1889). However, the cell phenotypes under selection in each organ remain unclear. Breast cancers are often categorized by molecular subtype for clinical purposes, defined by the presence of hormone receptors; these subtypes correlate modestly with preferential relapse sites, but still allow the possibility to metastasize to multiple sites (Soni et al. 2015; Kennecke et al. 2010; Smid et al. 2008; Cummings et al. 2014). Histological grade, defined by cell morphology, mitosis, and cellular differentiation state, also does not correlate well with tissue tropism (Cummings et al. 2014). The oncogenic mutations found in driver genes are also used to type breast cancers, but these mutations remain fairly consistent between primary tumors and untreated metastases (Reiter et al. 2018). It is therefore likely that metastasis tropism is determined by factors other than those used to type breast cancers at the clinical level.

Pre-clinical work has shown some of the other factors that command tropism, including cytokines and proteins secreted from tissues and tumor cells, the compositions of the immune microenvironment and oncogenic miRNAs (Obenauf and Massagué 2015). The metabolic preferences of cancer cells may also play a role in metastasis tropism (Dupuy et al. 2015). Tumor cells are long known to exhibit metabolic alterations (Warburg et al. 1927), and reprogrammed metabolism is even considered a “hallmark of cancer” (Hanahan and Weinberg 2011). Nevertheless, the role of metabolic alterations in metastasis tropism has arguably received less attention.

The MDA-MB 231 cell line was derived from a breast cancer patient and forms metastases in multiple organs in mice (Cailleau et al. 1974; Puchalapalli et al. 2016; El-Mabhouh et al. 2008). The mouse model was used to select *in vivo* for lineages that preferentially home to the bone, brain or lung (Bos et al. 2009; Minn, Gupta, et al. 2005; Kang et al. 2003). The result was a set of parental MDA-MB 231 cells and their matched derivatives which home to specific organs. A recent study used these cells to study transcriptomic alterations in micro-metastasis formed in the lung compared to the parental and brain-colonizing cells (Basnet et al. 2019). They found that mitochondrial electron transport Complex I, oxidative stress, and counteracting antioxidant programs were induced in pulmonary micrometastases, again suggesting a key role for metabolism in the adaptation of cancer cells when colonizing a distant organ. Another study compared the metabolomic profiles of brain- and bone-homing lineages with parental cells (Li et al. 2020). The lineages were all cultured in the same *in vitro* conditions, and the comparison revealed differences in intracellular metabolite levels, particularly in purine nucleotides. This study also found increased serine metabolism in all three metastatic lineages (brain, bone, and lung) compared to parental cells, and concluded that these pathways were necessary for metastatic cell growth.

Here, we used the MDA-MB 231 model to study metabolic fluxes in breast cancer cells with different tropism. Our analysis shows that understanding how different cell lineages utilize metabolic pathways differently may require more than transcriptomic and metabolomic profiling. We combined metabolomics with direct measurements of fluxes and used mathematical models constrained by the data to study the cells’ metabolic fluxes. We focused on the brain- (BrM2) and lung-homing lineages of MDA-MB 231; the lung and brain have distinct metabolic microenvironments that may contribute to metastatic selection (Zhang and Liu 2015; Parpura et al. 2014; McKeown 2014; Datta et al. 1980; Lottes et al. 2015). Our results show that the flux through the glycolytic pathway is faster in brain-homing and especially in lung-homing lineages compared to parental. Importantly, this occurs even though parental cells have higher intracellular levels of glucose and other intermediaries of the glycolysis pathway. The simulations carried out using our mathematical model show how cells can sustain a high glycolytic flux even though they have lower levels of glycolysis pathway intermediates: lung-homing cells in particular prevent feedback inhibition by constitutively activating a pathway sink into lactate, leading to elevated glycolysis activity. We confirmed this prediction by measuring glycolytic enzymatic activities directly, and we propose that the ratio of expression of lactate dehydrogenase genes to pyruvate dehydrogenase genes indicates a cell state of high glycolytic flux. We conclude that this ratio is clinically relevant by showing that it correlates with lung metastases in patient samples.

## Results

### Metabolomic and transcriptomic profiles show differences in glycolytic pathway in primary versus metastatic lineages

We profiled the metabolomes of MBA-MB 231 cell line and two of its metastatic derivatives, the brain-homing BrM2 lineage and the lung-homing lineage LM2 lineage (**Fig. 1A**). The profiles differed for each lineage, indicating that the lineages maintain heritable differences in their utilization of metabolic pathways even when cells they are cultured *ex vivo* in the same condition (**Fig. 1B**). Principal component analysis (PCA) of the metabolite levels shows that the largest differences occur between the parental line and the derived lineages (**Fig. 1C**): all replicates of the parental lineage scored high on PC1, which explained 84% of the variation. A biplot analysis revealed that a single metabolite—glucose—explained a large part of these differences (**Fig. 1C**, shown in gray). In fact, the relative glucose levels in parental cells were >30x higher than in either BrM2 or LM2 (BrM2 p=0.0004, LM2 p=0.001) (**Fig. 1C inset**). Most other intermediates of glycolysis were also significantly higher in the primary lineage, with the exception of higher 2,3-BPG and pyruvate in BrM2 relative to parental, and higher lactate in LM2 relative to parental (**Fig. 1D and Supplementary Fig. 1A**). The PCA also suggested that the primary metabolic divergence occurred early in the metastatic process, with further diversification in the lung and brain: PC2, which explained ~10% of the variation, distinguished BrM2 from LM2. The differences between BrM2 and LM2 occurred mostly in metabolites from amino acid and fatty acid pathways (**Supplementary Fig. 1B**).

**Figure 1:**
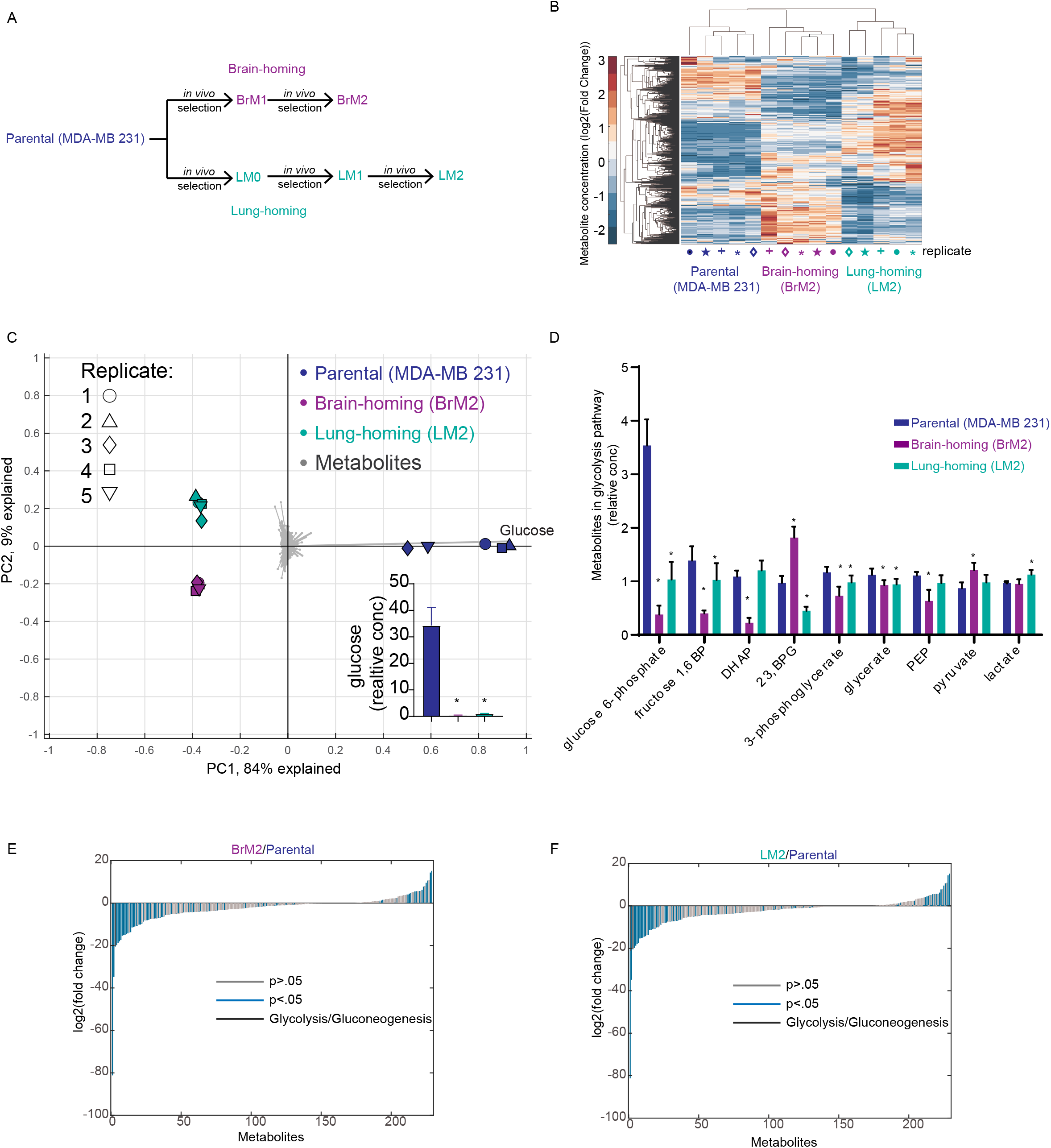
Integrated metabolomics and transcriptomics show differences in glycolysis pathway in primary versus metastatic lineages. **A)** Diagram of the *in vivo* selection, which started with the parental lineage MDA-MB 231 and made the brain-homing BrM2 and lung-homing LM2 derivatives. **B)** Heatmap of metabolite levels of parental and metastatic derivatives. Each cell line had 5 replicates which clustered together. **C)** PCA of parental, BrM2, and LM2 metabolite levels, overlaid with a biplot showing the correlations of individual metabolites. Inset: bar plot of glucose levels: fold-change relative to parental: BrM2: 73-fold, p=0.0004, LM2: 33-fold, p=0.001. Fold change LM2/BrM2: 2-fold, p=0.000007. Data are represented as mean ± SD. **D)** Barplot of metabolite levels of the other components in glycolysis. * indicates p<.05. Data are represented as mean ± SD. **E)** Waterfall plot of MITHrIL output comparing BrM2 to parental networks, highlighting the glycolysis/gluconeogenesis pathway. **F)** Waterfall plot of MITHrIL output comparing LM2 to parental networks, highlighting the glycolysis/gluconeogenesis pathway. **See also Figs S1 and S2**.

A closer look at the RNA expression profiles already published (Minn, Gupta, et al. 2005; Bos et al. 2009) showed that the expression of glycolytic pathway genes was also perturbed in the brain- and lung-homing lineages. Many of those genes were expressed at lower levels in the metastatic cells compared to parental cells, confirmed by enrichment of this gene set in parental cells using GSEA(Subramanian et al. 2005; Mootha et al. 2003) (**Supplementary Fig. 1C-D**).

We sought to integrate the transcriptomic and metabolomic data and investigate the pathways most affected throughout the entire metabolic network. For this we used MITHrIL (Mirna enrIched paTHway Impact anaLysis) (Alaimo et al. 2017)(Alaimo et al. 2016) (**Supplementary Fig. 2A**). MITHrIL confirmed that glycolysis was indeed among the most significantly perturbed pathways in both metastatic lineages relative to parental cells (**Fig. 1E-F and Supplementary Fig. 2B**). Overall, these results indicated that the components of the biochemical reactions in glycolysis were perturbed in the brain- and lung-homing lineages, and were largely lower than in parental cells. Of note, MITHrIL found that lactate metabolism was higher in LM2 but not BrM2 cells (**Supplementary Fig. 2C**). Further MITHril analysis comparing BrM2 to LM2 showed lower levels of brain-associated pathways in LM2, including synapse signaling, and higher oxidative phosphorylation and pyruvate metabolism compared to BrM2 cells (**Supplementary Fig. 2D**).

### Glycolysis flux and lactate secretion increased in metastatic lineages despite lower levels of glycolysis intermediates

We measured the rate of glucose uptake by the cells in balanced growth using a YSI analyzer. Interestingly, the glucose influx was not lower but marginally higher in the brain-homing lineage (1.3-fold, p=0.03) and markedly higher in the lung-homing lineage (2.0-fold, P=0.001) (**Fig. 2A**). Lactate secretion was approximately proportionally higher in those lineages: 1.1-fold in the brain-homing lineage (P=0.30) and 1.6-fold in the lung-homing lineage (P=0.002) (**Fig. 2A**).

**Figure 2:**
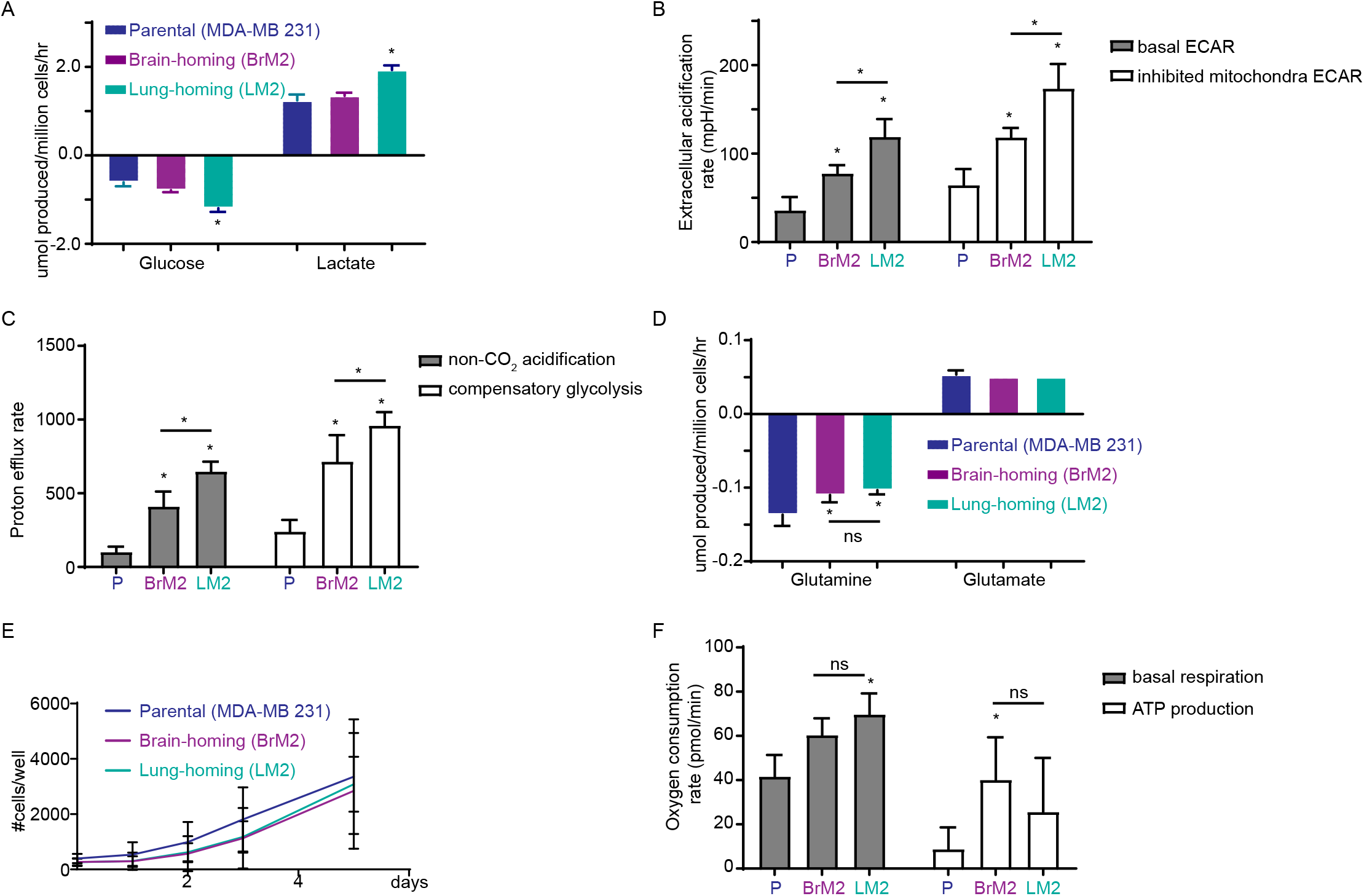
Glycolysis flux and lactate secretion increased in metastatic lineages despite lower levels of glycolysis intermediates. **A)** YSI analysis of glucose uptake and lactate production. Glucose fold-change relative to parental: BrM2: 1.3-fold, p=0.04, LM2: 2.0-fold, p=0.001. Fold change LM2/BrM2: 1.5-fold, p=0.002. Lactate fold-change relative to parental: BrM2: 1.1-fold, P=0.30, LM2: 1.6-fold, P=0.002. Fold change LM2/BrM2: 1.4-fold, p=0.001. **B)** Seahorse analysis of basal ECAR as well as ECAR after mitochondrial inhibition by antimycin A and rotenone. Basal ECAR fold-change relative to parental: BrM2: 2.2-fold, p=0.00002; LM2: 3.3-fold, p=0.0000004. Fold change LM2/BrM2: 1.5-fold, p=0.0003. Mitochondrial-inhibited foldchange relative to parental: BrM2: 1.8-fold, p=0.000009; LM2: 2.7-fold, p=0.0000005. Fold change LM2/BrM2: 1.5-fold, p=0.0002. **C)** Seahorse analysis of non-mitochondrial ECAR and compensatory glycolysis. Non-mitochondrial ECAR fold-change relative to parental: BrM2: 3.1-fold, p=2×10^-20^, LM2: 4.5-fold, P=9×10^-36^. Fold change LM2/BrM2: 1.4-fold, p=8×10^-12^. Compensatory glycolysis fold-change relative to parental: BrM2: 2.6-fold, p=4×10^-16^, LM2: 3.4-fold, P=1×10^-31^. Fold change LM2/BrM2: 1.3-fold, p=1×10^-8^. **D)** YSI analysis of glutamine uptake and glutamate production. Glutamine fold-change relative to parental: BrM2: .8-fold, p=0.11, LM2: .8-fold, P=0.04. Fold change LM2/BrM2: .9-fold, p=0.3. Glutamate fold-change relative to parental: BrM2: .9-fold, p=0.7, LM2: .9-fold, P=0.7. Fold change LM2/BrM2: 1-fold, p=1. **E)** Representative experiment showing growth rates of the three cell lineages. Final growth rates and fold changes were calculated using a generalized linear regression model on logged data from 4 independent experiments. Logged parental growth rate: .24 cells/day. Fold-changes: BrM2: .003-fold, p=0.93, LM2: −.0003-fold, p=0.99. When only the exponential phase of growth was considered, excluding lag-phase, BrM2: 1.05-fold, p=0.004, LM2: 1.03-fold, p=0.04. **F)** Seahorse analysis of basal respiration and ATP production. Respiration fold-change relative to parental: BrM2: 1.5-fold, p=0.001, LM2: 1.7-fold, P=0.00009. Fold change LM2/BrM2: 1.2-fold, p=0.06. ATP fold-change relative to parental: BrM2: 4.6-fold, p=0.002, LM2: 3.0-fold, P=0.1. Fold change LM2/BrM2: .6-fold, p=0.2. For all panels, data are represented as mean ± SD. **See also Fig S3.**

We then measured the rate of extracellular acidification using a Seahorse XF analyzer. The LM2 cells showed the fastest rates of extracellular acidification (BrM2: 2.2-fold, p=2×10^-5^; LM2: 3.3-fold, p=4×10^-7^) as well as acidification from non-mitochondrial sources (BrM2: 3.1-fold, p=2×10^-20^, LM2: 4.5-fold, P=9×10^-36^) (**Fig. 2B and C, left panels**). Inhibition of the electron transport chain with rotenone and antimycin A further increased extracellular acidification, with LM2 cells again releasing the highest levels of non-mitochondrial acidification among the three lineages (BrM2: 1.8-fold, p=9×10^-6^; LM2: 2.7-fold, p=5×10^-7^) (**Fig. 2B, right panel**). After this mitochondrial inhibition, adding 2-deoxyglucose halted glycolysis and led to a loss of extracellular acidification, indicating that the increase in acidification was indeed due to compensatory glycolysis in the absence of mitochondrial function (BrM2: 2.6-fold, p=4×10^-16^, LM2: 3.4-fold, P=1×10^-31^) (**Fig 2C, right panel**). Furthermore, YSI revealed that both derived lineages consumed slightly less glutamine than parental cells (BrM2: .8-fold, p=0.11; LM2: .8-fold, p=0.03), while glutamate secretion remained unchanged (**Fig. 2D**). It is therefore unlikely that glutamine underlies the faster rate of lactate production.

Our observations suggest that derived lineages—especially the lung-homing lineage— have an enhanced Warburg effect, the most prevalent purpose of which is thought to be for increased anabolism (Hitosugi and Chen 2014). Despite these differences, all three lineages grew at similar rates (BrM2: 0.003-fold, p=0.93, LM2: −0.0003-fold, p=0.99), suggesting that the metabolic adaptation we had observed served a different purpose than faster biomass production (**Fig. 2E**). Interestingly, when we considered only cells in the exponential phase of growth, and we excluded the initial lag-phase, the brain- and lung-homing lineages appeared to grow slightly faster than the parental lineage (BrM2: 1.05-fold, p=0.004, LM2: 1.03-fold, p=0.04). This suggests that established metastatic tumors may mildly outpace primary tumor cells in growth rate. However, the marginal increase in exponential-phase growth rate was smaller than the magnitude of metabolic changes observed above, giving more support to the notion that the faster influx of glucose serves another function (Liberti and Locasale 2016). We also confirmed that mitochondrial function was not impaired in the derived lineages: both lineages had higher mitochondrial respiration rates (BrM2: 1.5-fold, p=0.001, LM2: 1.7-fold, P=0.00009) and ATP production compared to parental, indicating that the glucose influx was also not required for glycolysis-derived energy in these cells (**Fig. 2F**). Overall, the mitochondrial metabolism of LM2 differed the most from that of parental, with BrM2 showing an intermediate level (**Supplementary Fig. 3**).

### Mathematical model and experimental validation explain higher flux despite fewer components

The data presented so far stress an important point: the levels of the metabolic intermediates of a pathway do not necessarily correlate with flux through that pathway. Brain- and lung-homing lineages have higher glucose consumption and lactate production than parental cells despite lower levels of the molecular components—intermediary metabolites and even the mRNAs of pathway enzymes—of the glycolytic pathway. These differences were marginal in brain-homing cells and robust in lung-homing cells. To understand how lung-homing cells may have these higher fluxes despite lower abundances of glycolysis pathway components we turned to a systems level analysis using mathematical modeling. We adapted a model of the flux rates in cancer cell metabolism (Balcarcel and Clark 2003): a 24-flux metabolic network model was constrained by the fluxes measured by YSI or Seahorse XF, while unknown fluxes were unconstrained. Given that the flux-balance solutions were not unique, we quantified the uncertainty of unmeasured fluxes by sampling the constrained high-dimensional flux space. Flux sampling allowed us to compute the most likely solution under the constraints given by data (**Fig. 3A-C and Supplementary Fig. 4A-C**).

Interestingly, the model suggested that metastatic lineages can flexibly modulate how nutrients are used for respiration/ATP production versus biomass. For example, flux from pyruvate to lactate was calculated to be approximately equal in parental and BrM2 cells, despite the higher glucose uptake in BrM2. BrM2 cells used the excess glucose-derived carbons for biomass and the TCA cycle while lowering the use of glutamine for either purpose (**Supplementary Fig. 4A**). This would allow BrM2 cells to “catch up” to the level of ATP production in LM2 cells despite lower glucose uptake than LM2 cells, while still maintaining similar rates of growth.

**Figure 3:**
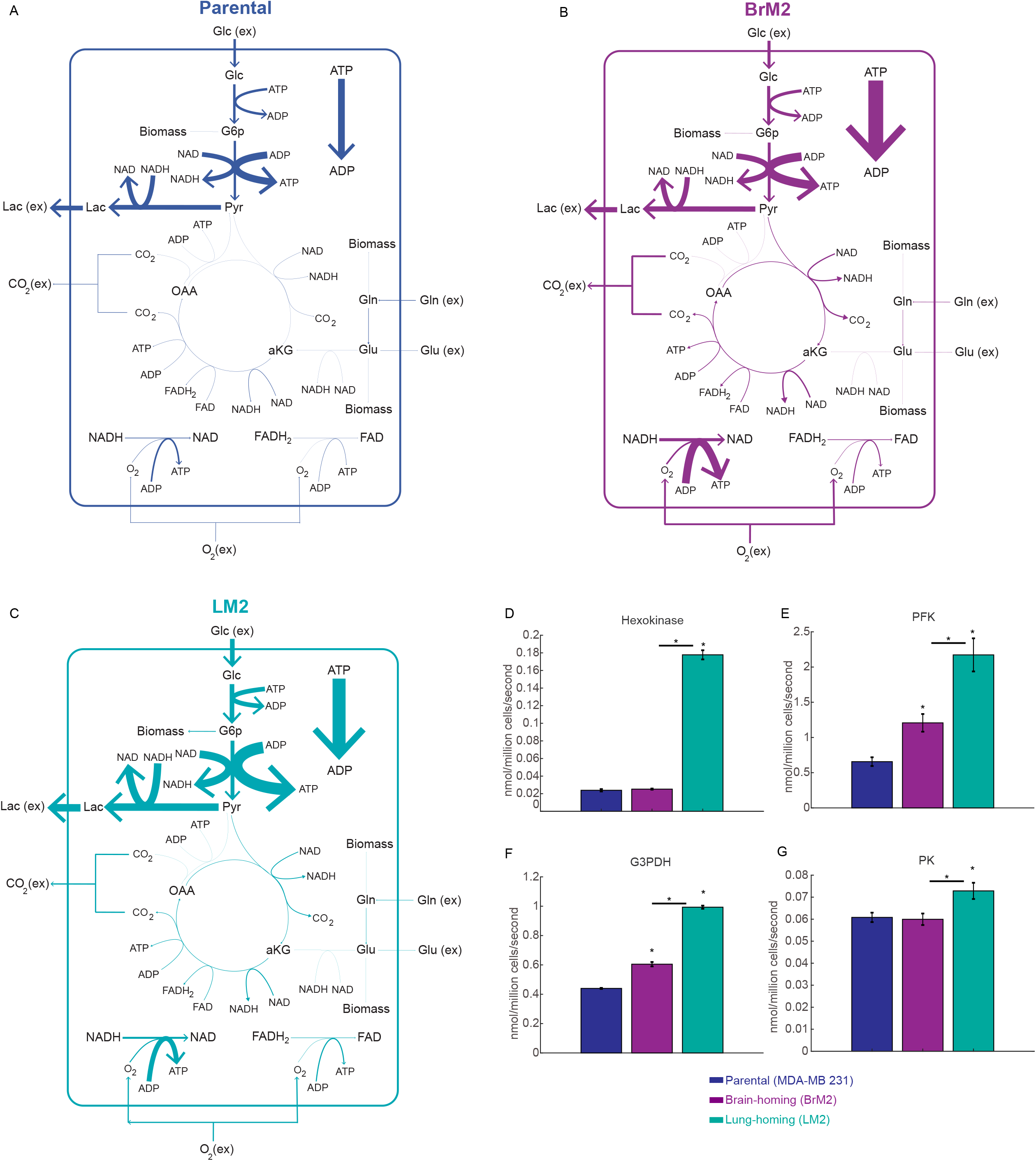
Mathematical model and experimental validation explain higher flux despite fewer components. **A-C)** Mathematical modeling of fluxes of select metabolic pathways for parental, BrM2, and LM2 lineages, respectively. Data from Figure 3 were used to constrain the model, and values for unknown fluxes were calculated. Fluxes are represented as most likely values. **D)** Enzymatic activity of hexokinase. Fold change relative to parental: BrM2: 1.1-fold, p=0.4, LM2: 7.4-fold, p=0.00007. Fold change LM2/BrM2: 7.0-fold, p=0.00006. **E)** Enzymatic activity of phosphofructokinase. Fold change relative to parental: BrM2: 1.8-fold, p=0.003, LM2: 3.3-fold, p=0.003. Fold change LM2/BrM2: 1.7-fold, p=0.004. **F)** Enzymatic activity of glyceraldehyde 3-phosphate dehydrogenase. Fold change relative to parental: BrM2: 1.4-fold, p=0.002, LM2: 2.3-fold, p=0.00001. Fold change LM2/BrM2: 1.6-fold, p=0.00003. **G)** Enzymatic activity of pyruvate kinase. Fold change relative to parental: BrM2: .98-fold, p=0.08, LM2: 1.2-fold, p=0.04. Fold change LM2/BrM2: 1.2-fold, p=0.03. For panels D-G, data are represented as mean ± SD. **See also Fig S4**.

These model predictions were confirmed by additional Seahorse measurements. The results showed that mitochondrial pyruvate utilization was higher in metastatic lineages compared to parental cells and mitochondrial glutamate utilization was lower in metastatic lineages compared to parental cells (**Supplementary Fig. 4D-E**).

Importantly, our model indicates that glycolytic flux should indeed be the highest in LM2 cells and lowest in parental cells (and that the high glucose uptake and lactate production in LM2 cells are not uncoupled). To confirm that parental cells perform less glycolytic flux despite higher metabolic intermediate levels, we directly tested the enzymatic activity of four key steps in glycolysis. The LM2 cells had indeed the highest levels of activity while parental cells had the lowest levels of activity (**Fig. 3D-G**), again confirming that the expression of glycolysis pathway genes (the mRNA levels) do not necessarily correlate with flux through that pathway.

### Elevated LDH and a high LDH/PDH ratio supports constitutive lactate efflux in lung metastases

Glycolysis is inhibited by its products (Berg et al. 2002). Therefore, in order to support a high flux it is necessary to maintain low levels of metabolic intermediates and prevent feedback inhibition. This can only be achieved by also sustaining flow into a “sink”: e.g. lactate efflux. In LM2 cells this correlated with an increased expression of the lactate dehydrogenase gene *LDHB* (**Fig. 4A-B**). Therefore, although counterintuitive, the lower levels of glycolytic intermediates in LM2 cells relative to parental are not in conflict with higher flux, but may be *required* to maintain high glycolysis flux in the absence of any other regulation. Pyruvate—the end product of glycolysis—can be converted either to lactate or to acetyl-coA. However, increased flux into acetyl-coA and subsequent mitochondrial activity increases ATP, and a high ATP/AMP ratio also allosterically inhibits enzymes in glycolysis (Berg et al. 2002). Therefore, directing pyruvate predominately to lactate rather than acetyl-coA could serve to maintain a high glycolytic flux. To determine whether this was indeed the case in our cell lines, we calculated the ratio of gene expression between lactate dehydrogenase and pyruvate dehydrogenase genes. Consistent with our model, LM2 cells had a significantly higher ratio of LDH/PDH genes than both parental and BrM2 lineages (**Fig. 4C**).

**Figure 4:**
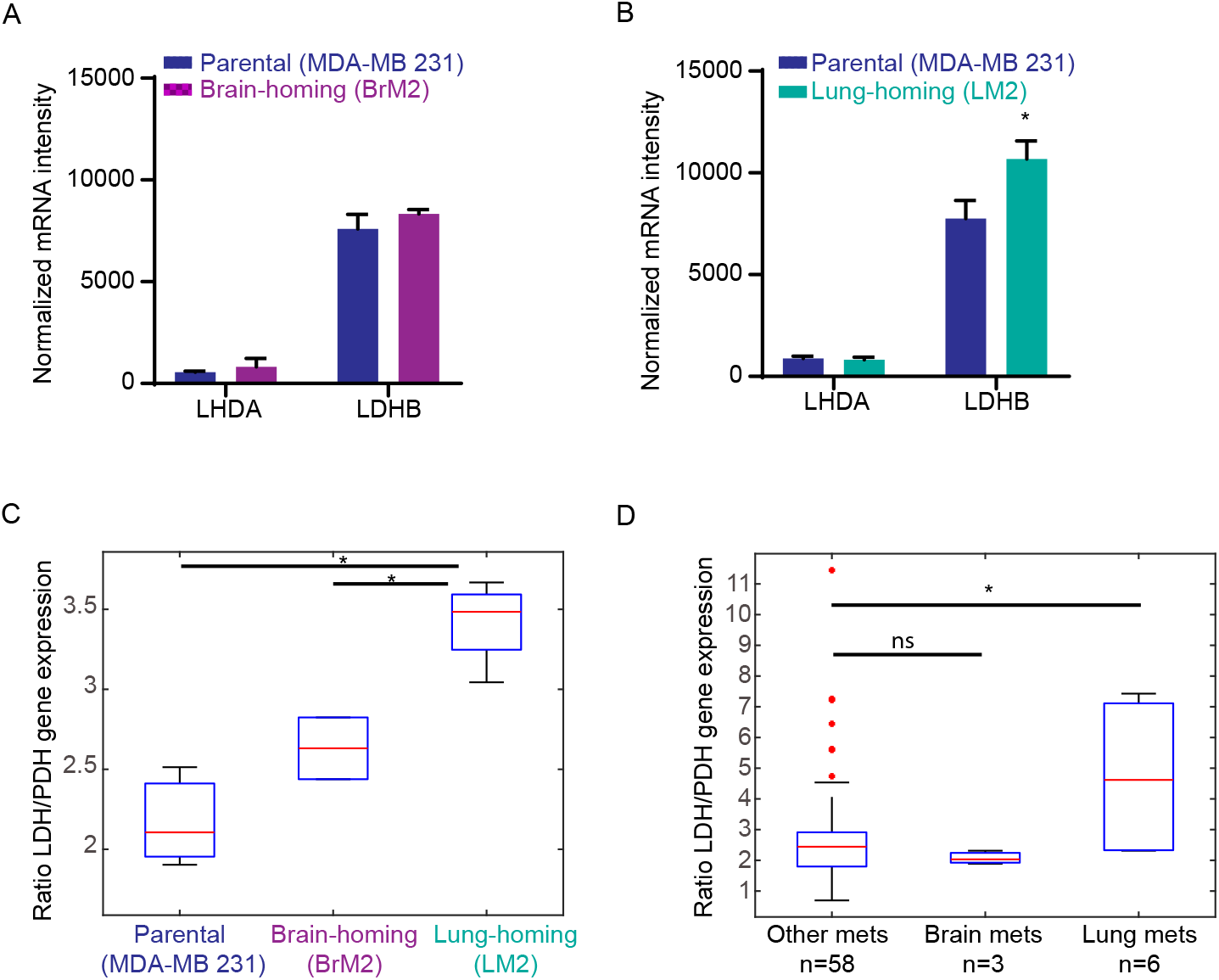
A high LDH/PDH ratio supports constitutive lactate efflux in lung metastases in both cell lines and clinical samples. **A)** RNA expression of LDHA and LDHB in BrM2 vs parental cells. LDHA: 1.5-fold, p=0.3, LDHB: 1.1-fold, p=0.3. Data are represented as mean ± SD. **B)** RNA expression of LDHA and LDHB in LM2 vs parental cells. LDHA: .9-fold, p=0.7. LDHB: 1.4-fold, p=0.02. Data are represented as mean ± SD. **C)** Boxplot of ratio of LDH genes (sum of LDHA and LDHB) to PDH genes (sum of PDHA1 and PDHA2) in cell lines. Fold change relative to parental: BrM2: 1.2-fold, p=0.2, LM2: 1.5-fold, p=0.007. Fold change LM2/BrM2: 1.3-fold, p=0.03. **D)** Boxplot of ratio of LDH genes to PDH genes in patient tumors from The Metastatic Breast Cancer Project. The “Lung” classification includes all tumors that had lung metastases, “Brain” classification includes all tumors that had brain/CNS metastases, and “Other” classification contains tumors that had metastases other than to the lung or brain/CNS. Fold change lung/other: 1.6-fold, p=0.02. **See also Fig S5.**

### Clinical data show that LDH/PDH gene expression signature is higher in breast cancers that metastasize to the lung

To test the clinical relevance of our findings, we asked whether the ratio of LDH/PDH expression was higher in the tumors of patient that developed lung metastasis. We analyzed the Metastatic Breast Cancer Project dataset (Wagle et al. 2016) which includes a diverse cohort of >100 patients with metastases in multiple sites. The most common site is bone, followed by liver, lymph node, lung, and brain/central nervous system (**Supplementary Fig. 5A**). In these data, as expected, the oncogenic lesions do not determine metastasis; this is clear in that the most commonly mutated genes, PIK3CA and TP53, (**Supplementary Fig. 5B**) do not correlate with metastatic site (**Supplementary Fig. 5C**). Also as expected, hormone receptor status does not correlate with metastasis (**Supplementary Fig. 5D**). In order to determine whether metastatic site correlated with transcriptomic state, we clustered the transcriptomes of all samples into six archetypes (**Supplementary Fig. 5E**). The top metastatic sites span several archetypes, indicating—also as expected—that the general transcriptomic signatures do not determine tropism (**Supplementary Fig. 5F**).

We then asked whether differences in the LDH/PDH ratio could determine tropism. We classified patients according to whether they had a lung metastasis, brain metastasis, or any metastasis other than to the lung or brain. Patients with lung metastases had a higher LDH/PDH ratio than patients with other metastases, supporting our hypothesis that this ratio characterizes a common adaptation in lung tropism (**Fig. 4D**). Patients with brain metastases did not have a higher LDH/PDH ratio than those with other metastases. Interestingly, the magnitude of the LDH/PDH ratio in patient samples did not cluster by archetypes identified by gene expression, indicating that the LDH/PDH ratio is independent of the major cell state in breast cancer (**Supplementary Fig. 5G**).

## Discussion

Our data, combining metabolomics, transcriptomic analyses, flux measurements, mathematical modeling and validation experiments lead to the following five conclusions: first, mRNA levels do not necessarily correlate with enzymatic activity; second, metabolic intermediates may *anticorrelate* with flux; third, different lineages evolved from the same line can have distinct heritable metabolic fluxes; fourth, metastatic lineages in our model display higher glycolytic flux and bioenergetics than parental cells, with lung-homing cells exhibiting by far the greatest glucose uptake and lactate production; fifth, a high LDH/PDH ratio maintains elevated glycolytic flux in lung-homing cells and in patients with lung metastases, which suggests that it is important for lung tropism.

Our mathematical model of fluxes was key to make sense of the data. Biomass production differed only slightly between the three cell lineages, and the model helped us understand how the derived lineages may increase or decrease uptake of nutrients in order to meet cellular demands. Taken together, our results suggest a demand-based model rather than a supply-based one. Rather than taking up all available nutrients at high rates to increase growth rate or energy output, cells may modulate uptake to meet specific needs: e.g. lactate production.

The metabolic alterations in brain-homing cells seemed modest compared to parental cells, especially when contrasted with the strong alterations we saw in lung-homing cells. This could indicate that the selection for metabolic adaptation was stronger in the lung. However, it could also indicate that all metastatic cells originally carried the adaptations found in LM2, but that the BrM2 lineage, in the process of overcoming additional challenges like crossing the blood-brain barrier, lost some of these metabolic alterations in favor of other, more necessary, adaptations to the brain microenvironment.

Metabolizing glucose by glycolysis to produce pyruvate and secrete lactate is energetically inefficient (Vander Heiden et al. 2009). Still, many cancer cells behave this way for reasons that may provide selective advantages beyond increased growth rates (Liberti and Locasale 2016). For example, lactate secreted can lower the pH of the microenvironment and may trigger tissue-repair responses in stromal cells that help tumor development (Carmona-Fontaine et al. 2017). Lactate has also been shown to increase migration and metastases by degrading the extracellular matrix (Bonuccelli et al. 2010). LM2 cells, and other breast cancer metastases to the lung, may be selected in part for their ability to produce lactate and thus form successful metastases in the lung. Interestingly, exogenous lactate decreases glucose utilization in the lung (Fisher and Dodia 1984). It is possible that this leads to more glucose availability for colonizing cells, leading to a feed-forward loop in lung metastases.

Gene signatures are typically sets of genes that are up- or down-regulated, with expression changes that may be independent of each other. Our results argue in support of gene signatures that are more complex (Itadani et al. 2008). When it comes to metabolism, a gene signature that reflects *balance* may be more functionally relevant than a set of genes that change concordantly. We found that the LDH/PDH expression ratio may be more important for the maintenance of high glycolytic flux, rather than the individual expression of either or even both genes. While there have been associations between LDH expression and the progression of different cancers (Mishra and Banerjee 2019), it will be interesting to see future cancer metabolism analyses incorporate the concept of metabolic balance and investigate metabolic genes that are interdependent. Interestingly, a mouse model of breast cancer metastasis found that metastatic cells can upregulate both glucose consumption/lactate production and oxidative phosphorylation compared to parental cells, consistent with our results, but that liver metastases regulate the balance between the two in a HIF-1α and hypoxia-dependent manner (Dupuy et al. 2015). This supports our idea that metabolic balance is more complex than the expression of genes taken independently, and that expression in the context of other genes and environmental cues is important.

The diverse metabolic changes we observed in the three lineages could be evidence that a primary tumor induced by a set of driver genes can still have underlying diversity at the metabolic level, driven by non-driver genetic differences. This would suggest that metabolic rewiring in cancer cells is more complex than single oncogenic changes (Levine and Puzio-Kuter 2010). However, we cannot rule out the possibility that the cells in the primary tumor that ultimately produced BrM2 and LM2 had mutations in driver genes that the rest of the primary tumor did not share, and were undetected in the primary tumor due to low abundance. Another possibility is that alterations in common driver genes occur at the transcript, rather than genomic level, in metastatic lineages. A recent study in MDA-MB 231 cell lines found overexpression of c-myc in metastases, especially in the bone-homing lineage (Li et al. 2020). Disseminated cells may also be metabolically plastic due to reversible epigenetic states that can be reprogrammed depending on the distal tissue (McDonald et al. 2017).

Perhaps most importantly, our work warns that metabolomic profiling alone—or even in combination with transcriptomic profiling—may not suffice to show how cells use their metabolic fluxes. The static pictures provided by metabolomics and transcriptomics may require a combination of flux measurements and mathematical models to show how cells utilize their metabolic fluxes. We focused on fluxes through central carbon metabolism, but we acknowledge that the metastatic process, including organ-specific metastasis, likely involves many other secreted and consumed metabolites, and we did not directly measure the flux for all relevant compounds (Lu et al. 2010). We also do not yet know whether the metabolic changes we identified were the cause of lung- or brain-specific metastasis or a byproduct of the selection at the distal organ, a question we will explore in future studies.

## Acknowledgements

We thank Joan Massague’s lab for providing all the cell lines used in this paper. We thank the Flow Cytometry core facility at MSKCC for sorting of our fluorescently-labeled cells. We also thank Justin Cross, Mirela Biresa, and Weige Qin from the Cell Metabolism core facility at MSKCC for help designing metabolic flux experiments, training on Seahorse equipment, and YSI analysis. Thank you also to members of Joao Xavier’s lab for input on this project. This work was supported by NIH Research Program Grants under award numbers R01CA229215 and U54 CA209975 to J.B.X. D.M. is also supported by the Alan and Sandra Gerry Metastasis and Tumor Ecosystems Center (GMTEC) Gerry Fellowship.

## Author Contributions

D.M. conducted experiments, analyzed data, contributed to mathematical modeling, and wrote the manuscript. J.X. revised the manuscript and supervised all experiments and analysis. C.L. designed and conducted the flux balance models. A.L.F., S.A., and A.F. conducted MITHrIL analysis.

---

**Supplementary Figure 1: A)** Heatmap of the glycolysis pathway metabolites, showing BrM2 vs parental and LM2 vs parental. **B)** (Top) Heatmap of the top 20% most different metabolites between BrM2 and LM2 cells, labeled by pathway. (Bottom) Table of the top 5 metabolic pathways different in BrM2 vs LM2 cells. **C**) Heatmap of RNA expression of the genes in the Hallmark Glycolysis gene set, showing BrM2 vs parental and LM2 vs parental. **D)** Gene Set Enrichment Analysis of the Hallmark Glycolysis gene set, plotting enrichment scores for parental relative to LM2 (top) and parental relative to BrM2 (bottom).

**Supplementary Figure 2: A)** Diagram of MITHrIL algorithm. **B)** Heatmap of significantly differently regulated pathways determined by MITHrIL for BrM2/parental and LM2/parental cells. **C)** MITHrIL output for the glycolysis/gluconeogenesis pathway, overlaid on Kegg diagram of the pathway. Nodes (metabolites) and connections (genes) are colored according to strength of MITHrIL prediction. Top: BrM2/parental, bottom: LM2/parental. **D)** Heatmap of significantly differently regulated pathways determined by MITHrIL for LM2/BrM2 cells.

**Supplementary Figure 3:** PCA of the oxygen consumption rates of the three lineages under various mitochondrial inhibitors.

**Supplementary Figure 4: A)** Relative fold change of each flux for BrM2/parental (left) and LM2/parental (right). **B)** Distribution of fold flux frequency of BrM2/parental (left) and LM2/parental (right). Bars of the histogram represent the frequency that a given flux value was selected as fit to the model over many possible samplings; vertical lines indicate the most frequent (and therefore most likely) flux ratio. **C)** Distribution of flux level versus frequency for each flux in the model over many possible samplings. **D)** Seahorse analysis of pyruvate utilization in mitochondria. Fold change relative to parental: BrM2: 1.4-fold, p=0.4, LM2: 3.6-fold, p=0.02. Fold change LM2/BrM2: 2.6-fold, p=0.06. Data are represented as mean ± SD. **E)** Seahorse analysis of glutamine utilization in mitochondria. Fold change relative to parental: BrM2: .3-fold, p=0.02, LM2: .5-fold, p=0.3. Fold change LM2/BrM2: 1.9-fold, p=0.5. Data are represented as mean ± SD.

**Supplementary Figure 5: A)** MBCP analysis of the most common metastatic sites found in breast cancer patients. **B)** MBCP analysis of the most common mutations found in breast cancer patients. **C)** Stacked barplot showing the distribution of the most common metastatic sites relative to mutation status in MBCP patients. **D)** Stacked barplot showing the distribution of the most common metastatic sites relative to hormone receptor status in MBCP patients. **E)** UMAP (McInnes et al. 2018)of transcriptomics of MBCP patients (plus six dummy samples corresponding to each archetype center), colored by which archetype they best fit. Samples cluster by archetype but archetypes also overlap, indicating shared transcriptomics. “X”s mark the archetype centers, representing the mock samples that would perfectly fit each archetype. **F)** Stacked barplot showing the distribution of the most common metastatic sites relative to transcriptomic archetype in MBCP patients. **G)** UMAP of transcriptomics of MBCP patients, colored by LDH/PDH ratio. Patients with lung metastases are circled.

## STAR Methods

### All codes and raw data files are available on github

https://github.com/dm2791/Divergent-use-of-metabolic-fluxes-in-breast-cancer-metastasis

### Cell culture

All cell lines were grown in DMEM (Fisher 11965118) supplemented with 10% FBS (made in the MSKCC media core facility) and 1% penn/strep (Fisher 15140122). Cells were grown in a 37°C incubator with humidity and 5% CO_2_. Authenticated cell lines were obtained from the Massague lab and generated as described previously (Minn, Gupta, et al. 2005; Bos et al. 2009).

### Cell growth assay

Cell lines were infected with H2B-YFP or H2B-mcherry lentivirus using 20□g/mL polybrene. After a few days of passage, cells were sorted via Fluorescence-activated cell sorting. Cells were counted and plated in equal numbers in a 96-well plate, and fluorescent images were taken periodically for several days on a Zeiss microscope. A 5x objective was used in the instrument, and images were either collected once every 24h after which the plate was returned to the incubator, or once every hour using the custom-made chamber surrounding the microscope that maintained temperature, CO_2_, and humidity levels. Images were analyzed using the Zeiss Zen blue software and our own custom-made MATLAB scripts. Data from several independent biological replicates of the growth assay were pooled, and analyzed using a generalized linear mixed-effects regression model with a log function (since the cells are in exponential growth) and random effects for which experiment and well in the plate the data came from. Growth rates were analyzed both for the full data as well as excluding pre-exponential phase growth.

### Metabolomics

Cells were seeded in T75 flasks, in 5 replicates per cell line. 2 days after plating, cells were trypsinized, counted, and spun down. Media was aspirated from cells, which were then resuspended in PBS and re-spun. PBS was aspirated and the cell pellet was immediately frozen in liquid nitrogen. Samples were shipped to Metabolon on dry ice, and metabolite abundance data was normalized to total protein content per sample. Metabolomics analysis: heatmaps, principal component analysis, statistical analysis of abundance fold changes (two-sample *t*-test), and partial least squares regression analysis were done in MATLAB using built-in features and custom-made scripts.

### RNA expression analysis

mRNA expression data that were previously published (Bos et al. 2009; Minn, Gupta, et al. 2005) were downloaded from Gene Expression Omnibus. The Hallmark Glycolysis gene set was downloaded from the Gene Set Enrichment Analysis Molecular Signatures Database. Expression fold change relative to parental for each metastatic lineage was calculated for the genes in the Hallmark Glycolysis gene set, and a heatmap was generated using MATLAB. The ratio of (LDHA+LDHB)/(PDHA1+PDHA2) for each lineage and statistical differences (two-sample *t*-test) were calculated in MATLAB.

### MITHrIL pathway analysis

The MITHrIL algorithm was used as described previously (Alaimo et al. 2016). We used the combined Log2FC values of the differentially expressed genes (DEGs) and altered metabolites identified from BrM2 and LM2 samples. Specifically, the DEGs for BrM2 and LM2 samples were identified from the two public microarray projects used above while the altered metabolites were identified from our metabolomics data. Microarray data were first normalized and then the DEGs were identified by using the LIMMA package (Bioconductor) (Ritchie et al. 2015). Only the genes with Log2FC > 0.6 or Log2FC < −0.6 with a statistically significant adjusted p-value (with Benjamini-Hochberg correction) < 0.05 were considered differentially expressed and selected for the MITHrIL pathway analysis. The metabolomics data were analyzed using the MetaboDiff package (Bioconductor) (Mock et al. 2018) and only the metabolites with Log2FC > 0.6 or Log2FC < −0.6 and a statistically significant adjusted p-value (Benjamini-Hochberg correction) < 0.05 were considered as altered metabolites and included in the MITHrIL pathway analysis. All these analyses were performed using the framework Rstudio (R 3.5.2). MITHrIL output for the glycolysis/gluconeogenesis pathway was overlaid on the KEGG pathway diagram. Total MITHrIL output was imported into MATLAB to generate waterfall plots and clustergrams.

### YSI

Cells were cultured in 6-well plates. 8 hours prior to sample collection, media was changed to 2mL per well, and incubated as above. 1.2mL were collected from each sample, spun down to remove cell debris, and then frozen until processing by the Cell Metabolism Core Facility. Statistical analysis of data was done in MATLAB (two-sample *t*-test).

### Seahorse

20,000 cells/well were seeded in Seahorse XF plates. The following day, media was changed to the Seahorse XF media with glucose, pyruvate, and glutamine, and standard Seahorse XF protocols were followed. For assessing mitochondrial activity, the Mito Stress Test kit was used, with 1□M oligomycin, 1□M FCCP, and .5□M rotenone+antimycin A. Basal respiration rate corresponded to oxygen consumption excluding non-mitochondrial oxygen consumption (OCR in the presence of rotenone + antimycin A). ATP production corresponded to the difference in basal respiration rate and oxygen consumption in the presence of an ATP synthase inhibitor (oligomycin). For assessing glycolysis, the Glycolytic Rate kit was used, with .5□M rotenone+antimycin A and 50mM 2-deoxyglucose. Non-CO_2_ ECAR was calculated by subtracting mitochondrial OCR*.6(standard scaling factor) from the total proton efflux rate. Compensatory glycolysis corresponded to proton efflux rate in the presence of rotenone+antimycin A. For assessing mitochondrial fuel oxidation, the Mito Fuel Flex Test kit was used with 6 M BPTES and 4□M UK5099. Mitochondrial glutamine utilization was calculated by subtracting OCR in the presence of BPTES from total OCR. Mitochondrial pyruvate utilization was calculated by subtracting OCR in the presence of UK5099 from total OCR. All Seahorse data were normalized to cell count, and statistical analyses of fold change differences were done in MATLAB (two-sample *t*-test).

### Mathematical modeling

A 24-flux metabolic network model that represents glycolysis, reduction of pyruvate to lactate, TCA cycle, glutamine/glutamate metabolism, and oxidative phosphorylation was developed by simplifying the model reported in (Balcarcel and Clark 2003). Experimentally measured glucose uptake, lactate production, glutamine uptake, glutamate production, and oxygen consumption rates (converted to mmol/gDW/hr by assuming averaged cell weight of 1 ng) were used to constrain the corresponding fluxes by setting their lower and upper bounds to the mean measured value minus and plus standard error across all replicates respectively. Other fluxes were unconstrained with their lower or upper bounds set to −100 or 100 respectively. Considering the flux-balance solutions are not unique, we quantified the uncertainty of unmeasured fluxes by sampling the constrained high-dimensional flux space. Flux sampling allows building marginal distributions of each flux and computing the averaged flux value as the most possible solution under the constraints given by data. For the distribution of flux ratios between metastatic derivatives and parental cell types, we used a bootstrap method to resample the marginal distributions with replacement and calculate the fold change of resampled fluxes. Custom Python codes were developed with the COBRApy package (Ebrahim et al. 2013) to carry out all metabolic flux modeling and simulations in the paper.

### Enzymatic assays

The Hexokinase Activity Assay Kit (Abcam ab136957), Phosphofructokinase Activity Assay Kit (Abcam ab155898), Glyceraldehyde 3 Phosphate Dehydrogenase Activity Assay Kit (Abcam ab204732), and Pyruvate Kinase Assay Kit (Abcam ab83432) were used and accompanying protocols were followed. Briefly, cells from each lineage were trypsinized, counted, and 250,000 cells were collected for each assay. Cells were washed with PBS, homogenized, and the supernatant was mixed with substrate and probe developer. Absorbance was read at 450nm, and activity rate was calculated by measuring absorbance at two time points and compared to a standard curve. Negative and positive controls were used.

### Metastatic Breast Cancer Project analysis

The results included here include the use of data from The Metastatic Breast Cancer Project (https://www.mbcproject.org/), a project of Count Me In (https://joincountmein.org/) [(Wagle et al. 2016)]. Clinical metadata, genomic data, and gene expression data for the Metastatic Breast Cancer Project was downloaded from the cBio portal (MBCproject cBioPortal data version March 2020) (Cerami et al. 2012; Gao et al. 2013). Custom MATLAB scripts were used to calculate the most common metastatic sites and mutations in the selected samples. Archetypes were generated by using non negative matrix factorization, with each sample having probabilistic membership into each archetype. The maximum likelihood archetype was assigned as the given archetype for each sample. Proportions of each metastatic site for each breast cancer mutation, hormone receptor status, and transcriptional archetype were calculated in MATLAB. Patients were then classified according to whether they had metastases to the lung (including patients who had metastases to other sites as well as lung) or metastases to any location other than lung. Patients who had no metastases (or did not have any annotation regarding metastasis) were excluded. The ratio of gene expression of (LDHA+LDHB)/(PDHA1+PDHA2) was calculated for each patient; boxplot and statistical analysis of fold change between groups (two-sample *t*-test) was done in MATLAB. UMAP (McInnes et al. 2018) analysis of patient archetypes was done in MATLAB after z-scoring each sample.

