## Supplementary Figures for "Divergent use of metabolic fluxes in breast cancer metastasis"

Supplementary Figure 1

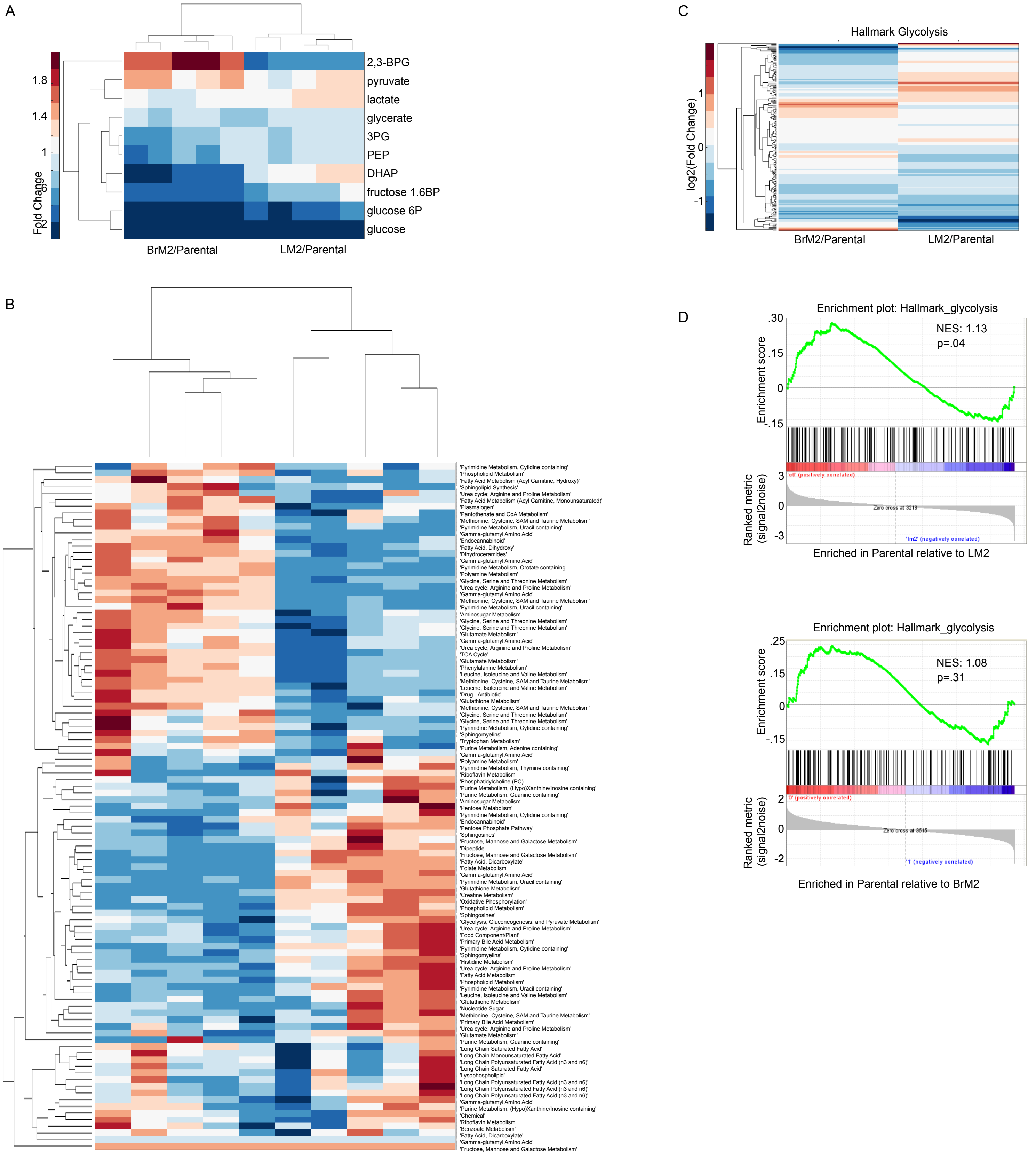


| | |
|---|---|
| Top pathways up in LM2 relative to BrM2 | 'Gamma-glutamyl Amino Acid'<br>'Pyrimidine Metabolism, Cytidine containing'<br>'Pyrimidine Metabolism, Uracil containing'<br>'Glycine, Serine and Threonine Metabolism'<br>'Purine Metabolism, Guanine containing' |
| Top pathways down in LM2 relative to BrM2 | 'Methionine, Cysteine, SAM and Taurine Metabolism'<br>'Urea cycle; Arginine and Proline Metabolism'<br>'Glutathione Metabolism'<br>'Long Chain Polyunsaturated Fatty Acid (n3 and n6)'<br>'Long Chain Saturated Fatty Acid' |

Supplementary Figure 2

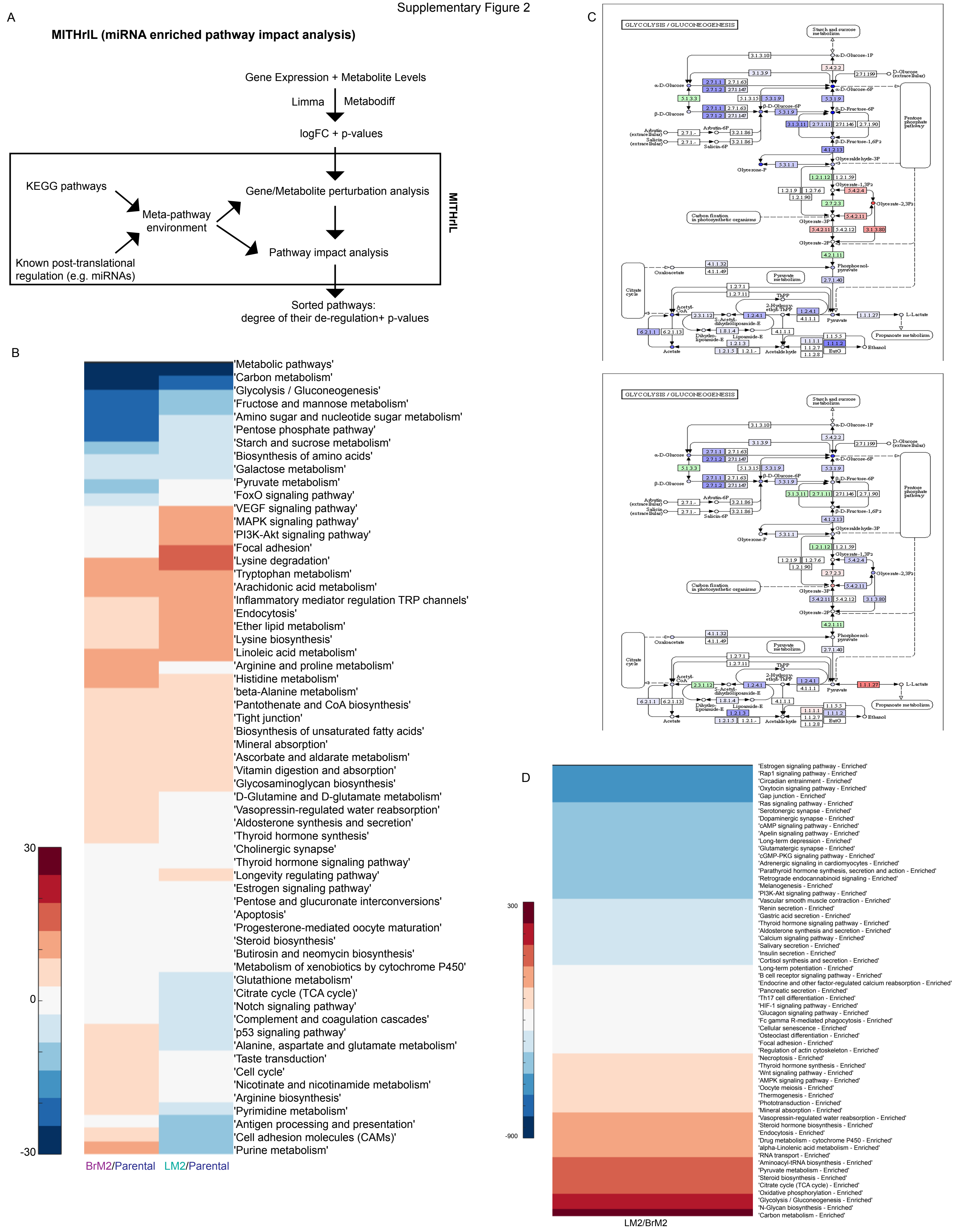

Supplementary Figure 3

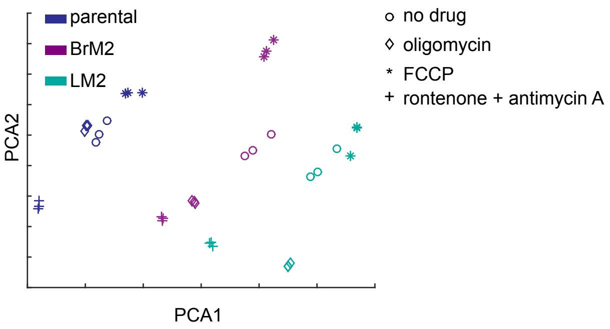

Supplementary Figure 4

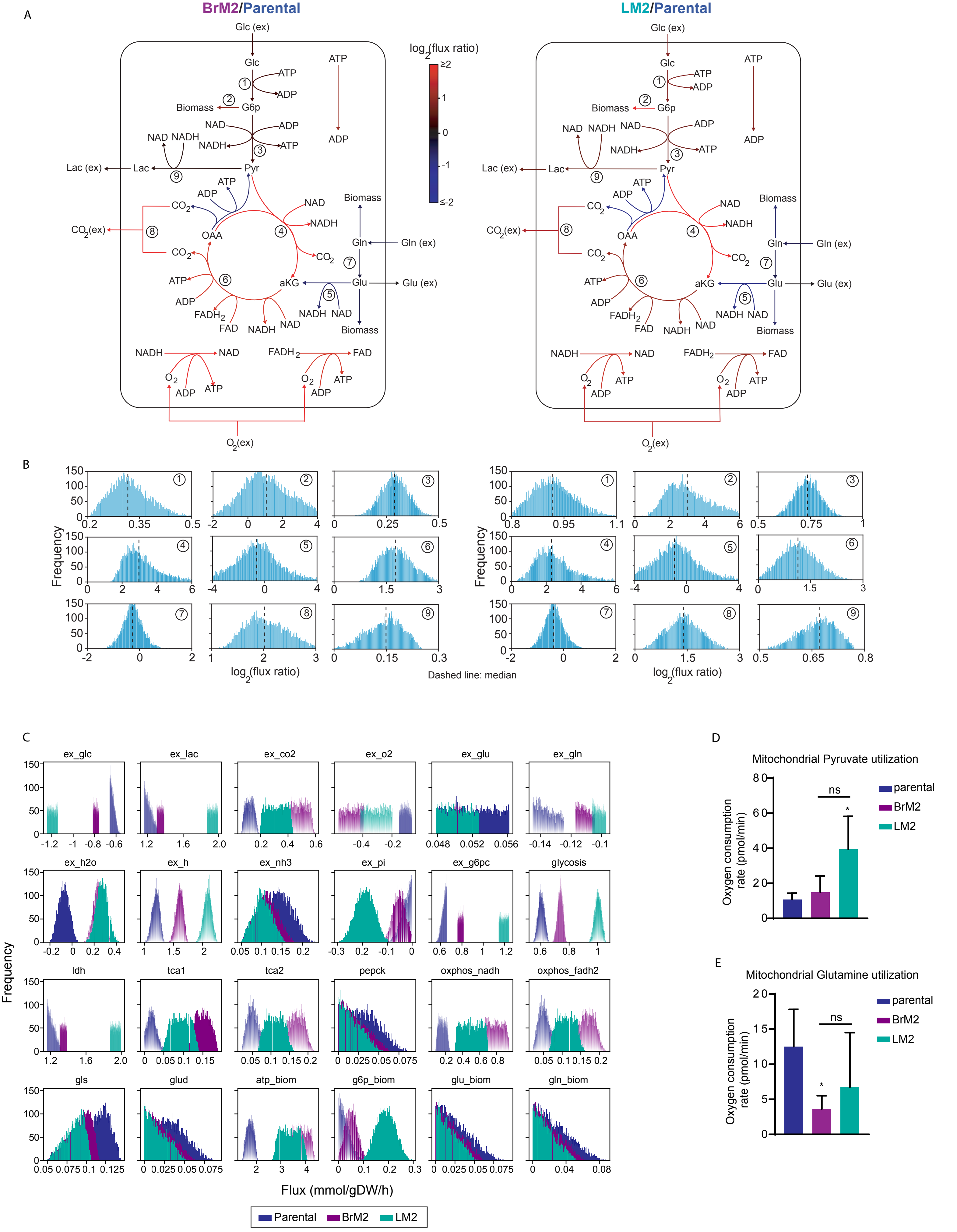

Supplementary Figure 5

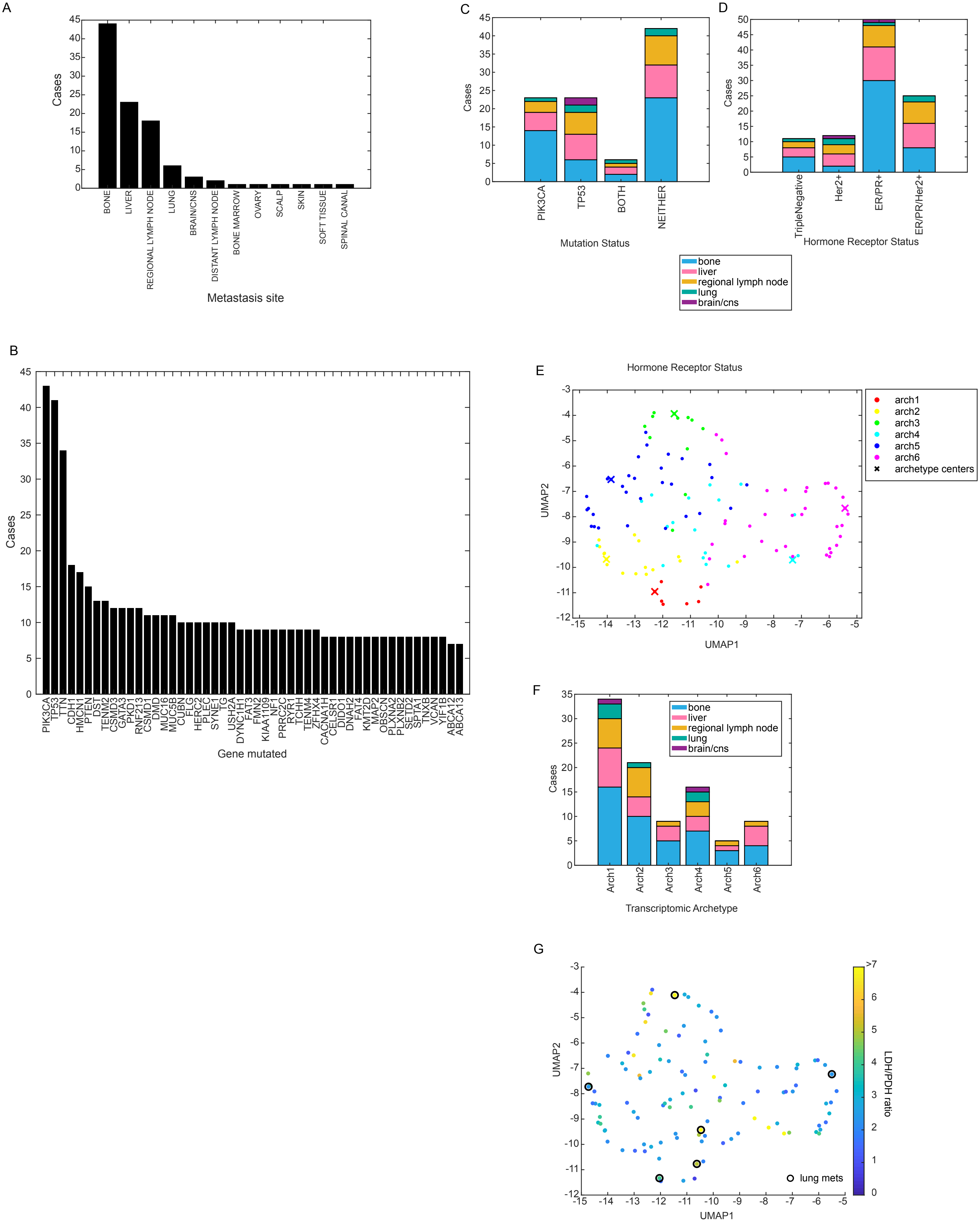
